# Biomarkers for aging identified in cross-sectional studies tend to be non-causative

**DOI:** 10.1101/624270

**Authors:** Paul G. Nelson, Daniel E. L. Promislow, Joanna Masel

**Author notes:** Corresponding author: Paul G. Nelson.

## Abstract

Biomarkers are important tools for diagnosis, prognosis, and identification of the causal factors of physiological conditions. Biomarkers are typically identified by correlating biological measurements with the status of a condition in a sample of subjects. Cross-sectional studies sample subjects at a single timepoint, while longitudinal studies follow a cohort through time. Identifying biomarkers of aging is subject to unique challenges. Individuals who age faster have intrinsically higher mortality rates and so are preferentially lost over time, in a phenomenon known as cohort selection. In this paper, we use simulations to show that cohort selection biases cross-sectional analysis away from identifying causal loci of aging, to the point where cross-sectional studies are less likely to identify loci that cause aging than if loci had been chosen at random. We go on to show this bias can be corrected by incorporating correlates of mortality identified from longitudinal studies, allowing cross sectional studies to effectively identify the causal factors of aging.

---

Despite the universality of aging (1, 2), we do not know which factors are the most important contributors to age-associated decline in physiological function and to increase in mortality rate. Further, we lack consensus regarding what it means to be aged and how to properly measure aging (3, 4). The advent of high throughput “omics” technologies has led to the tantalizing promise that a large enough dataset of informative biomarkers might both elucidate the causal factors behind aging, and provide precise, individualized diagnoses (5). Biomarkers have also been proposed as surrogate endpoints, allowing studies to forgo the slow and expensive process of measuring the symptoms of aging as they develop over time (6). However, the limits of what biomarkers can tell us about aging, and the consequences of the methods by which we discover biomarkers, have yet to be examined theoretically.

Biomarkers, broadly speaking, are biological quantities that are convenient to measure, and that give us medically important information (7, 8). Biomarkers can inform three very different categories of medically important information. First, biomarkers can inform diagnosis and associated prognosis. The first molecular biomarker was glucose, used to diagnose diabetes (9). Historically, sweet urine indicated death within a matter of months (10), making glucose in the urine useful for informing prognosis but little else. Today, prognosis without treatment is still useful for patients with untreatable cancer or Alzheimer’s disease, who may wish to put their affairs in order. Biomarkers for aging can similarly predict lifespan remaining or future quality of life (e.g. 11).

Second, biomarkers can reveal disease mechanisms. Perhaps surprisingly, while glucose is important to the mechanism of diabetes, historically, it did not lead physicians directly to the causal mechanisms of the disease. When medieval physicians noticed that glucose in the urine varied with diet, they mistakenly concluded that diabetes must be a disease of the digestive system (10). Instead, the indirect advantage of the discovery of glucose was that because its presence in the urine aided diagnosis, it facilitated the search for other mechanistic clues, specifically abnormalities in the pancreas noted during autopsy (12). In contrast, after cholesterol was noted as a prognostic biomarker for coronary artery disease, this led more directly to understanding the mechanistic role of plaque build-up (13).

Third, a biomarker can provide information that enhances interventions and improves outcomes (14). Such markers are called “predictive” rather than the “prognostic” biomarkers discussed above (15, 16). Insulin treatment, combined with a more accurate blood test (17), made glucose an important biomarker to drive treatment decisions (10). The presence of hormone receptors in breast cancer biopsy tissue is another predictive biomarker used to make treatment decisions. There is hope that the many cancer driver mutations currently being identified might also move from prognostic indicators to drug targets, in the way that oncofusion protein BCR-Abl (18) is targeted by Gleevec.

Identifying biomarkers generally begins with noting a correlation between a measurement and patient fate (19–22). Patient fate can be measured longitudinally, or else a known correlate of patient fate can be measured within a contemporaneous cross-section. To measure biomarkers of aging in a longitudinal study, potential markers are measured at the start, and a cohort is then followed through time to determine mortality. The correlation between marker status at the beginning of the study and later mortality rates or other symptoms of aging is then ascertained (23–25). In a cross-sectional study, biomarkers are identified based on their ability to predict either current chronological age (26), or a composite of chronological age and physiological indicators often termed “biological age” (27), even when their intent, as described above, is to predict lifespan remaining (28). New technologies mean that we can now investigate many potential biomarkers simultaneously, which opens the door both to discovering more and/or better markers, and to the risk of spurious results (29).

Longitudinal studies are obviously slow and difficult to conduct, but cross-sectional studies may not yield reliable results. Individuals with lower intrinsic mortality rates are likely to survive to older ages than their peers with high mortality rates, and are thereby more likely to be observed at older ages. This bias, known as cohort selection (30), may complicate the search for biomarkers of aging. Here, we simulate core aspects of the search for epigenetic biomarkers in the cross-sectional approaches, specifically those of Horvath and collaborators (26, 27, 31). We include the possibility of cohort selection, in order to determine what cross-sectionally identified biomarkers can, and cannot, tell us about aging.

## Methods

We first conduct simulations based on the procedure of Horvath (26), who trained an algorithm to predict chronological age from DNA methylation status at 21,389 sites in a cross-section of 7,844 individuals of different ages. The resulting regression model used 353 of those sites, and was used to assess the relative rate of aging (assessed as the difference between predicted age and chronological age) in a wide range of test datasets, from cancer tissue to patients with progeria.

Guided by this procedure, we simulate the methylation status of many loci during aging, followed by cross-sectional approaches to select loci as biomarkers. Specifically, we simulate *L*=20,000 loci in each individual. Each locus has two states, degraded and non-degraded, and begins the simulation in the undegraded state. We simulate enough individuals to obtain 1000 living individuals at each target age of 10, 20, 30,…80 years, with simulations proceeding in discrete time steps of one year. Each year, each un-degraded locus *i* has a probability *ρ_i_* of degrading, making the mean age at which the locus degrades (1-*ρ_i_*)/ *ρ_i_*.

We consider two models for the probability of dying *m*. First, we make the probability of dying increase exponentially with time, independently of which loci have degraded, according to a Gompertz mortality curve (32):

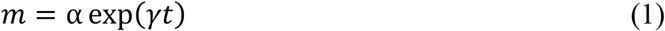

Using mortality rates from Arias et al. (33), we estimate *α* = *e*^−10^ deaths/year and *γ* = 0.0807/year to obtain a survival curve that approximates human demography in a developed nation, excluding elevated infant and early adult mortality, as well as any late life mortality deceleration.

In the second model, we make mortality a function of the number of degraded loci, choosing a function that yields a mortality curve that increases approximately exponentially with age. Specifically, we make mortality a power function of the sum of the effects of all degraded loci:

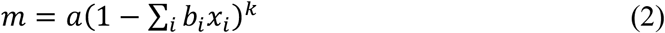

where *b_i_* is the effect size of locus *i* (often 0) and *x_i_* is 0 if the locus is in an undegraded state and 1 if the locus is degraded. Log mortality then has a linear relationship with age (i.e. we have a Gompertz mortality curve) in the special case where Σ*_i_ b_i_* = 1, with *k* determining its slope. With *k* < 0, mortality varies from a minimum of *a* when no causal sites have degraded to a maximum of infinity (instant death) when all sites have degraded. In our simulations, we set *b_i_* for all causative sites to equal 1/*L* where *L* is the number of causative loci. We refer to the “effect size” as the percent increase in mortality resulting from a single degraded locus in an otherwise undegraded individual: 100 × ((1 − *b_i_*)^*k*^ − 1).

To parameterize Equation 2 to approximate the Gompertz curve parameter values of Equation (1), we assume (just for the purpose of this parameterization) that all loci degrade at the same rate *ρ*, and thus the expected number of degraded loci as a function of time *t* is E(Σ_*i*_ *x_i_*) = *L*(1 − (1 − *ρ*)^*t*^). If we ignore variation both in effect sizes of causative loci and in the number of causative loci degraded, and we also ignore cohort selection (i.e. ignoring the fact that individuals with more than average degradations are less likely to be alive to have their mortality assessed), substitution of this expectation into Equation 2 would yield:

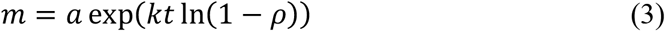

We therefore set *a* = α and *k* = *γ*/ln(1 − *ρ*) in Equation 2. Figure 1 shows that when these simplifying assumptions are relaxed, mortality in our Equation (2) simulations (red and blue for small and large *b_i_*, respectively) still exhibits the exponential relationship with time characteristic of a Gompertz mortality curve (solid black), with a slight deviation toward late life due to cohort selection when loci are of large effect (blue).

**Figure 1:**
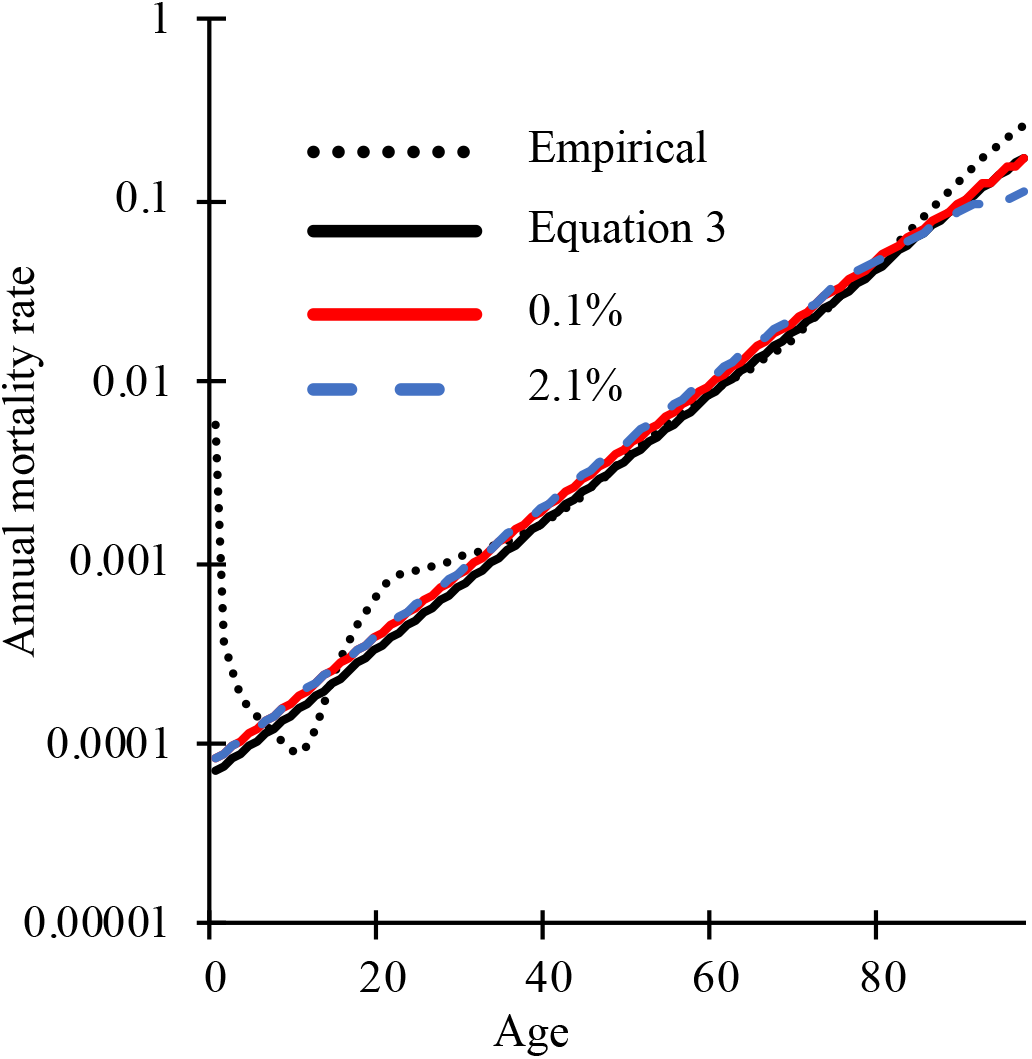
The Gompertz mortality curve (solid black) approximates the empirical estimates of human mortality (dotted black), excluding increased infant and early adult mortality (33). To include cohort selection, Equation 2 was simulated for a starting population of 1000 individuals using 20,000 loci of which 6,251 loci have an effect size of 0.1% (red) or 301 loci have an effect size of 2.1% (blue).

Some more recent efforts to identify biomarkers of aging (e.g. Levine (27)) train on a measure of “biological age” that incorporates phenotypic indicators in addition to chronological age. Phenotypic indicators are physiological measurements identified by their ability to predict, within a linear regression, mortality rates measured in a longitudinal study.

To determine how including measures of biological age affects the search for causal biomarkers of aging, we construct a simulated “biological age” phenotype *p*. We assume that phenotype can be scaled such that the mean phenotype of each age group is equal to the mean mortality rate of that age group (i.e., 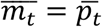), and that once this is done, the variance in phenotype among individuals of age *t* is equal to the variance in mortality (i.e., var_*t*_(*m_i_*) = var_*t*_(*p_i_*)). We also assume that within any age cohort, the phenotype correlates with mortality with the same correlation coefficient *r_p,m_*. Finally, we assume *p* is normally distributed, making individual *i*’s simulated phenotype:

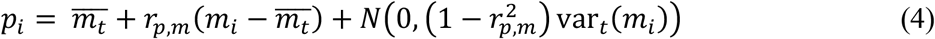

where *m_i_* is the true mortality rate of individual *i*, and *N*(0, *σ*^2^) is a random number drawn from a normal distribution with mean 0 and variance *σ*^2^. This formulation lets *r_p,m_* determine the fraction of variance in *p* that stems from variation in *m*, without changing var_*t*_(*p_i_*).

Next we consider how phenotypes can be used to construct a biological age. Simulated phenotypic values could be used, along with mean mortality 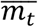 for each age group, to predict individual mortality rates using a linear model:

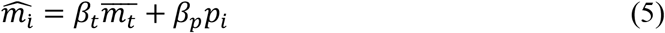

Instead of simulating the fit of this regression model to a data sample, we consider the best possible predictor by obtaining values for *β* analytically when the number of datapoints *n* goes to infinity. First, we take the expectation of both sides of Eq. 5, giving 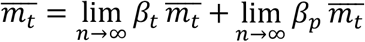, hence for the best predictor we will have *β_t_* = 1 − *β_p_*.

Second, we minimize the sum of squares 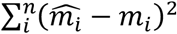. From Equation 5,

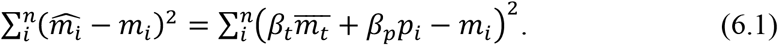

Using *β_t_* = 1 − *β_p_* yields:

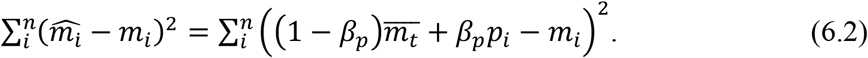

Substituting in Equation 4 yields:

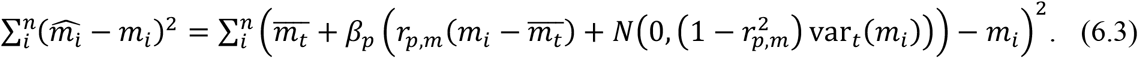

We next take the limit as the number of data points goes to infinity. Using

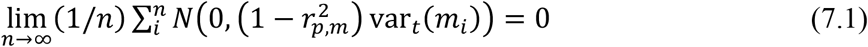

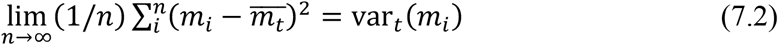

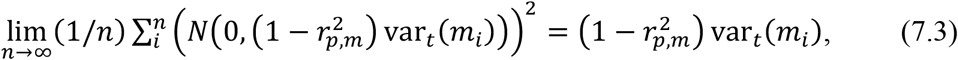

we obtain, after some rearrangement,

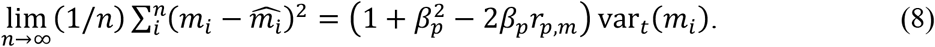

This reaches a minimum when the derivative by *β_p_* is zero, which occurs when *β_p_* = *r_p,m_*.

To obtain an estimated biological age 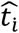 from the combination of an observed phenotype *p_i_* and a chronological age *t*, using this idealized linear model, we back-transform the estimated mortality from Equation 5 using the Gompertz mortality function (Equation 1):

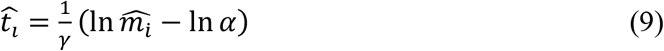

For each age cohort, we simulate each individual until they either die or reach the target age; only the latter are included in the training dataset. The process is repeated until we obtain 1000 individuals for each of the 8 ages of interest. Each locus is coded as 1 if degraded at the age of sampling and 0 otherwise. We correlate loci status either with chronological age, as in Horvath (26), or with biological age, as in Levine et al. (27), using the glmnet package in R (34). Regression coefficients between locus status and age are generated with a ridge lasso elastic net with the elastic net mixing parameter α=0.5 as in Horvath (26). During our analysis we noticed subtle biases in the regression coefficients generated by glmnet as a function of the order in which the loci appear in the training data set. To prevent such biases from affecting our results, we randomized the order of loci in the data set.

## Results

First, we illustrate the effect of cohort selection on the degradation status of age-causing and neutral loci over time (Figure 2). Let *n_i_* be the number of individuals with *i* degraded loci, *m_i_*(*t*) the probability of dying from age *t* to age *t*+1 of individuals with *l* degraded loci, and *p_binom_*(*l − i, L − l, ρ_i_*) the probability from a binomial distribution of *l − i* loci degrading out of *L − l* previously non-degraded loci when the probability of a locus degrading per year is *p_i_*. The expected number of individuals of age *t* years with *l* degraded loci is then:

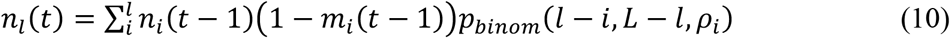

**Figure 2:**
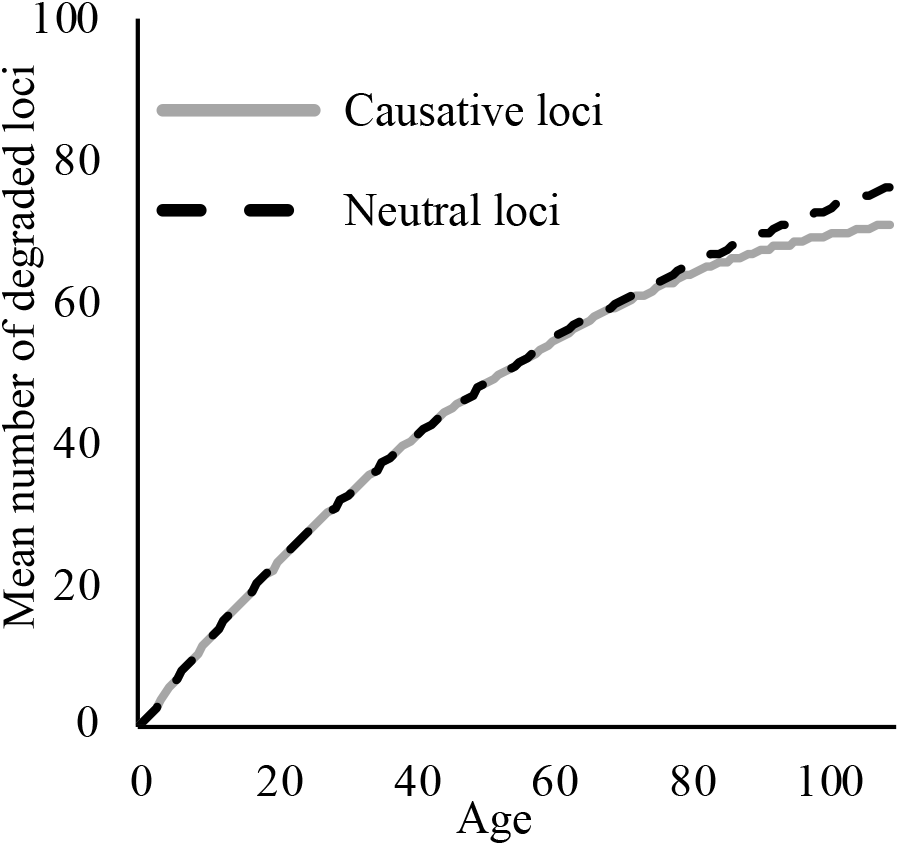
The mean number of degraded loci per individual when loci cause aging (grey), or are neutral (dashed black) out of 301 total loci. We use the largest effect size shown in Figure 3, where the degradation of one causative locus results in a 2.3% increase in mortality rates in an otherwise non-degraded individual, and all loci have an expected age of degradation of 75 years.

For neutral loci, we make mortality a function of time as given by Equation 1 (Figure 2, dashed black); for causative loci, mortality is given by Equation 2 (Figure 2, solid grey). Early in life, the frequency of degraded age-causing loci is dominated by the rate of degradation and is thus indistinguishable from neutral loci. Later in life, the additional mortality incurred by age-causing loci results in a slight decline in frequency relative to non-causal loci. We will see below that even though cohort selection is subtle as illustrated in Figure 2, it is nevertheless enough to make loci that cause aging slightly worse correlates of chronological age than neutral loci.

Next, we examine the effect of two locus attributes – degradation rate, and the effect size of degradation on mortality – on the regression coefficient assigned by glmnet as a predictor of age. Loci with expected ages of degradation less than 40 make poor biomarkers of age, and the most commonly selected biomarkers have expected ages of degradation between 60 and 100 (Figure 3A). Similarly, the most informative loci (i.e. those assigned the largest weights by glmnet) are expected to degrade between ages 70 through 90 (Figure 3B).

**Figure 3:**
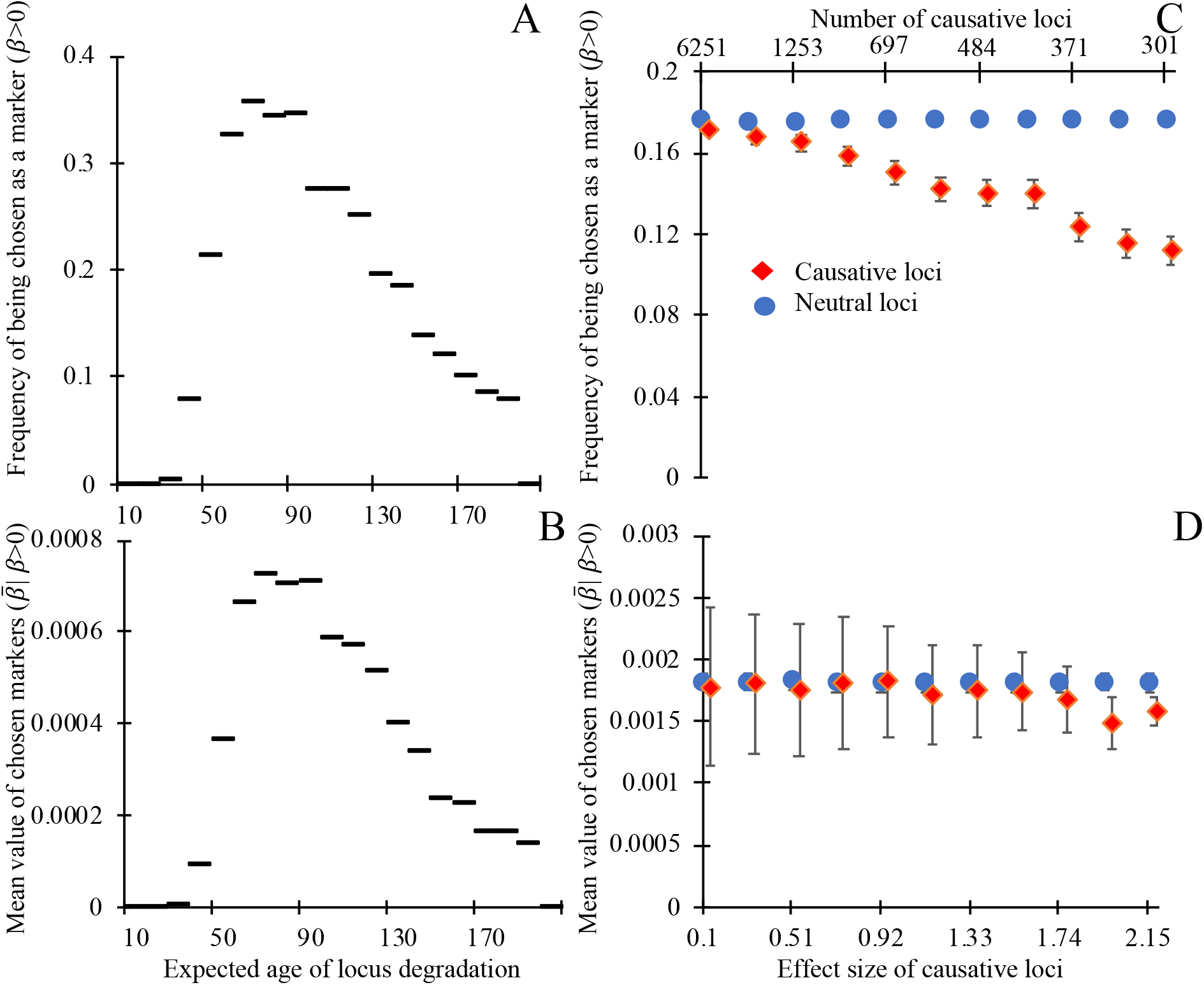
Biomarkers selected on their ability to predict chronological age tend not to be causative for aging, and degrade, on average, late in life. Loci whose expected time of degradation is late in life (age 50+) are chosen as biomarkers of aging (A) more often and have larger regression coefficients (in units of degradation status/year) when chosen as biomarkers (B), than loci that degrade earlier in life (note that all loci in panels A and B are non-causative). Causative loci (red diamonds) are less likely to be selected as biomarkers (C) than neutral loci in the same genome (blue circles). Of the loci selected as biomarkers (D), the magnitude of regression coefficients of causative loci is slightly smaller, on average, than those of neutral loci, at least when causative loci are of fairly large effect (>2%). Panels A and B show the outcome of a single simulation in which each of the 20,000 loci have expected ages of degradation 1+199(*i*/20,000) for integers *i* = 0 through 20,000. All loci are neutral and mortality is determined by Equation (1). Horizontal lines indicate the y-axis value corresponding to the 200 loci in each bin. A degradation time well above human lifespans indicates a locus with a low probability of degrading prior to death. Each point in panels C and D shows the mean outcomes of six independent replicate simulations in which each of the 20,000 loci are either neutral or have a single causal effect size (non-round-number effect sizes are an artifact of making choices using an alternative effect size metric). Mortality is determined by Equation (2), with the numbers of causative loci (top x-axis C,D) inversely proportional to the effect size of a causal locus (bottom x-axis C,D). In C and D, all loci have an expected age of degradation of 75 years. Error bars show standard errors; markers for causative loci have been offset slightly to the right.

Loci that have no causal effect on mortality are more likely to be chosen as informative markers (Figure 3C) than their age-causing counterparts. This counter-intuitive result arises because, as shown in Figure 2, cohort selection makes age-causing loci correlate more weakly with age than neutral loci. This biases regression analyses trained on age toward neutral markers. In the rare cases when age-causing loci of large effect are chosen, they have slightly smaller regression coefficients than neutral loci (Figure 3D). Together, these results show that loci that cause aging are worse predictors of chronological age than neutral loci.

Training on chronological age biases regression analysis away from identifying causative loci of aging, but more recent work instead trains biomarkers on “biological age,” a combination of chronological age and phenotypic measurements that are known to correlate with mortality (27). Such phenotypic measurements are identified by regressing phenotype and mortality over the course of a longitudinal study (e.g. 24).

To determine whether training on longitudinally validated correlates of aging can help identify causative loci of aging in a cross-sectional study, we modified our analysis to incorporate chronological age, a known correlate of individual mortality, and simulated noise.

We construct a simulated biological age-revealing phenotype such that the correlation coefficient *r_p,m_* between an appropriately transformed version of that phenotype and mortality has the same value within each age cohort. Biological age is then calculated as an optimal function of phenotype and chronological age (see Methods). When 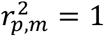, biological age perfectly reflects an individual’s true mortality; when 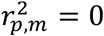, it gives no information beyond that already given by chronological age. Figure 4 shows that when phenotype provides little additional information 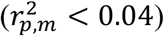, non-causative loci (blue circles) are preferentially chosen as biomarkers to predict biological age. However, when phenotype is a reliable indicator of mortality 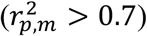 most, or even all, causative loci (red diamonds) are chosen as biomarkers of age.

**Figure 4:**
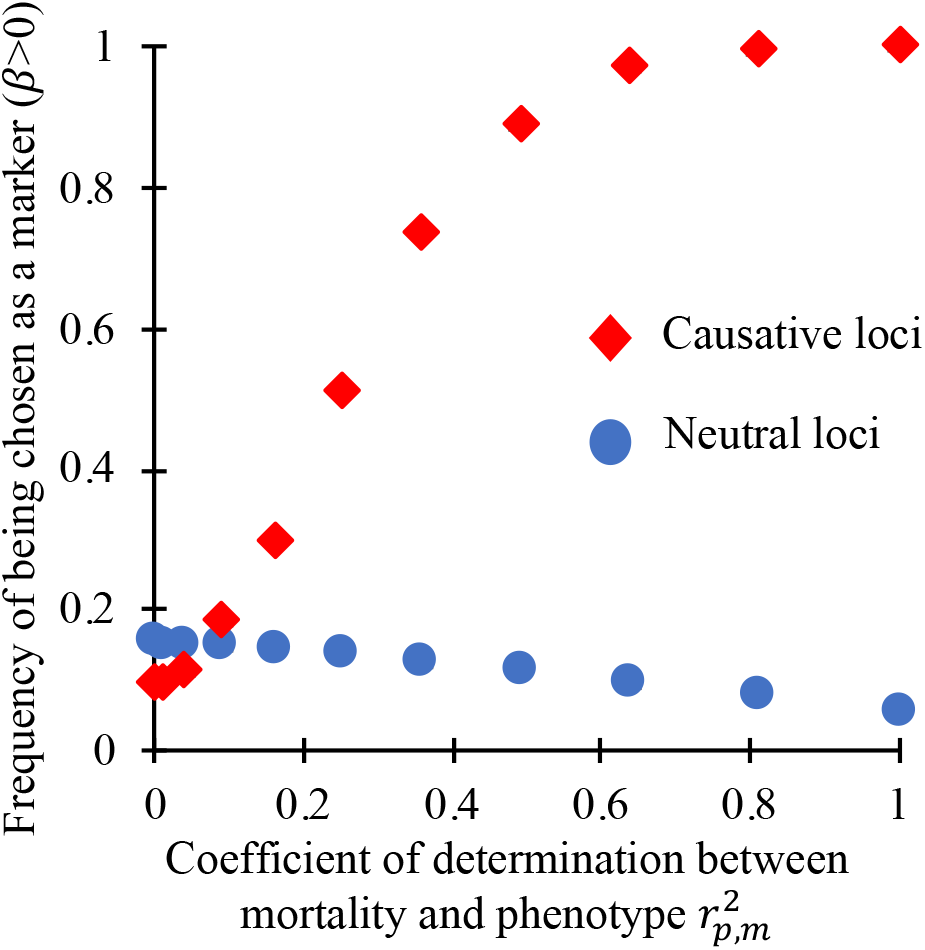
When phenotypes can accurately distinguish individual mortality within a cohort of the same age, regression on biological age (here an optimal function of both chronological age and phenotype) can preferentially choose loci that cause aging (red diamonds) over neutral loci (blue circles) as biomarkers of aging. As in the rightmost markers in Figure 3 C and D, here 301 loci of large effect are causative of aging.

To evaluate the quality of existing age-revealing phenotypes in the context of the search for causative biomarkers of aging, we used NHANES III linked laboratory and mortality data from the CDC (35, 36). We regressed (using the lm function in R) the nine phenotypic indicators used in Levine et al. (27) (white blood cell count, serum alkaline phosphatase, red cell width distribution, mean blood cell volume, lymphocytes percent, ln(serum C-reactive protein), serum creatinine, serum albumin, and serum glucose) on lifespan remaining, including only the 6898 individuals who had died over the course of the study. Chronological age explained *r*^2^ =0.184 of variation in lifespan remaining, while the combination of chronological age plus phenotype explained *r*^2^ =0.28, indicating that the amount of unique information provided by phenotype is *r*^2^ = 0.096 with respect to lifespan remaining, on the cusp of where biological age is useful for identifying causative loci of aging in our idealized model with respect to mortality rate.

However, there is also considerable stochastic variation in lifespan remaining even when there is no variance in underlying mortality rate. It therefore seems clear that the phenotypic age markers used by Levine et al. (27) are above the threshold quality needed to identify causal loci of aging more often than chance.

Next, we examine whether biomarkers of aging may be useful prognostic indicators of variation in the overall rate of degradation between individuals. To test this, we generated populations where the rate of aging varies among individuals, and populations where individuals vary in initial status, but then continue to age at the same rate. We trained the model to predict chronological age from loci status, as above. Comparing the difference between predicted age and chronological age to the known underlying rate of aging shows that biomarkers, even neutral biomarkers, can provide useful information regarding the rate of aging (Figure 5A), but not regarding initial mortality rate (Figure 5B). Note that we examined the special case in which variation in either the slope or intercept of the aging curve arises at birth. These results should still hold when it is later events that alter longevity, e.g. following an intervention. Biomarkers of aging could therefore inform the efficacy of interventions that affect the slope of the Gompertz curve, but not its intercept.

**Figure 5:**
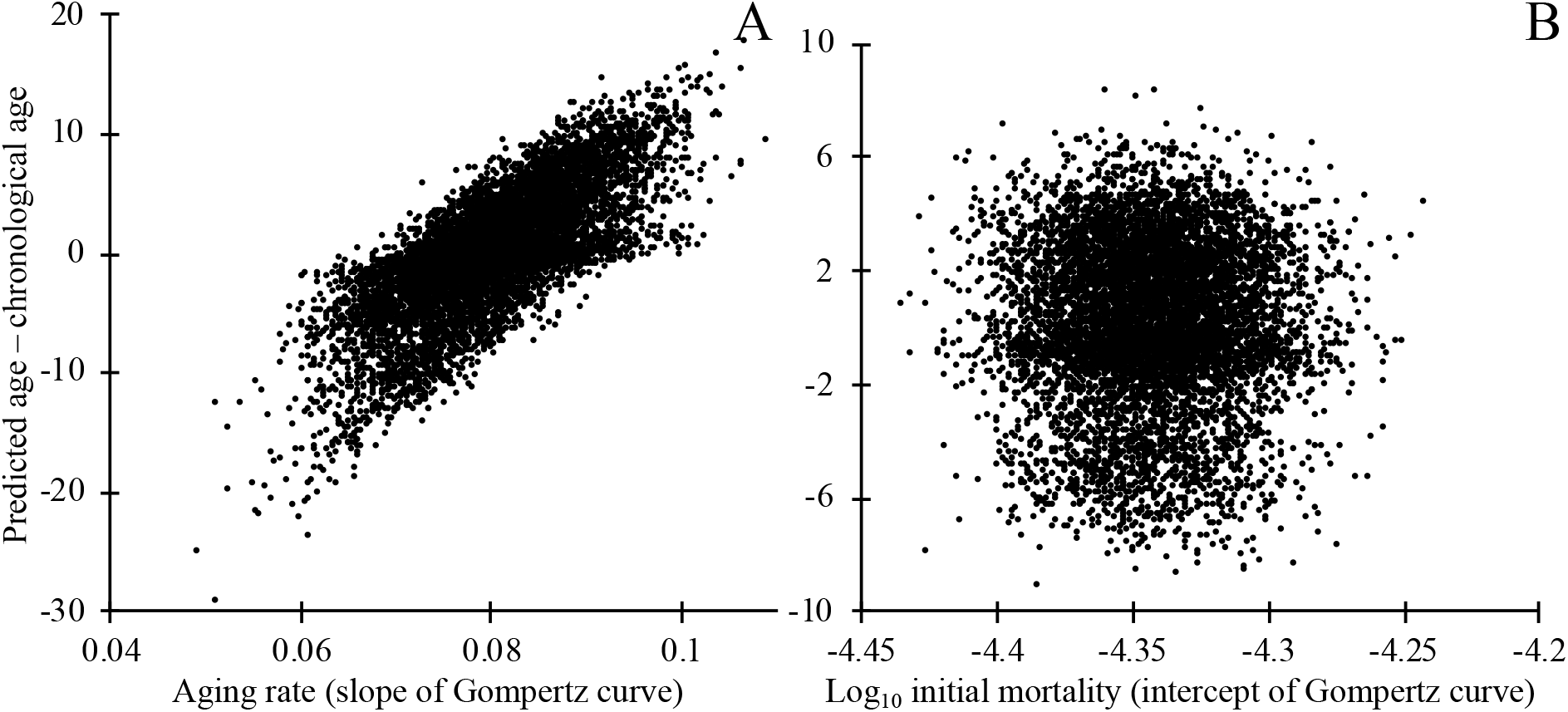
The deviation between an individual’s chronological age and predicted age informs the rate of aging, as measured by the slope of the Gompertz mortality curve (A), but not the intercept of the Gompertz mortality curve (B). Mortality depends on causal biomarker loci according to Equation 2. Markers were chosen by regression on chronological age in a training dataset, then applied to a separate testing dataset of independent but identically constructed simulations. In A, each individual’s rate of aging was drawn at birth from a normal distribution with mean 0.0807 and standard deviation 0.00807. The rate of aging *γ_i_* of individual i determines the probability of degradation *ρ_i_* of that individuals’ loci, where *ρ_i_* = 1 − exp(*γ_i_* ln(1 − *ρ*) /0.0807) where *ρ*=1/76, which gives an estimated age of degradation of 75 years as in Figure 3; all loci are non-degraded at birth. In B, a number of loci drawn from a Poisson distribution with mean 10,000 were degraded at birth, making the mean mortality rate at birth of individuals in the top quartile roughly 17% higher than those of the bottom quartile. To keep the mean mortality rate at birth *m_0_*=*e*^−10^ as above, we make *a*=6.65×10^−7^; the rate of degradation of loci is constant (*ρ*=1/76) across all individuals. In both A and B, 301 out of 20,000 loci are causative of aging.

## Discussion

Like other biomarkers discussed in the Introduction, biomarkers of aging can have three uses: to estimate lifespan remaining (prognosis), to identify the mechanisms that cause aging (37, 38), and, most ambitiously and usually late in their development, to distinguish classes of individuals for whom different treatments are appropriate. In cross-sectional studies, epigenetic biomarkers of aging have been selected for their ability to predict age at the time of sampling (21). However, the individuals in the training dataset are not random samples from their age cohort. Specifically, some individuals will not be sampled because they died before reaching the age of interest, and these individuals will on average be faster-aging. The resulting bias is known as cohort selection. Here, we have shown that cohort selection can make a cross-sectional study design spectacularly ineffective at identifying causal factors of aging.

While we have focused on cohort selection associated with causal locus-induced mortality, a similar sampling bias can arise from study exclusion criteria. Sicker individuals might be less likely to enroll in such studies, and may be excluded, e.g. if they have defined diseases whose prevalence increases with age, such as cardiovascular disease or diabetes (39). In a cross-sectional study, any alleles or epigenetic events that cause a predisposition toward such diseases will thus be underrepresented in older individuals.

Even if cross-sectional analyses trained on chronological age cannot identify loci that contribute to accelerating mortality, such analyses are not without value. Much effort has been put into quantifying natural variation in the slope and intercept of the aging curve (40) as well as the impact of interventions (41, 42). Our simulations show that aging biomarkers can provide prognostic (and hence potentially diagnostic) information regarding variation in the overall rate of degradation among individuals (25), but not in the basal mortality rate. They may therefore be useful in identifying factors affecting the overall rate of aging, as suggested by Horvath (26).

Biomarker selection enriches not just for non-causal loci, but also for loci with an expected age of degradation toward the end of human lifespan. To obtain this result, we considered the degradation rate of a locus to be independent of its causal effect. Whether selection against age-related mortality, which is necessarily subtle (43), can lead to lower degradation rates in age-causing loci than in the rest of the genome is an important theoretical and empirical question. Degradation rates are a function of evolvable factors such as sequence context, chromatin structure, or enzyme recruitment (44–46). The evolution of site-specific error rates has been observed for species with larger population sizes than humans (47–49). Comparing methylation changes associated with the expression of genes known *a priori* to play a role in aging to those in putatively neutral sites may help illuminate whether loci that cause aging are under tighter regulation than the genome as a whole.

While we have focused on molecular biomarkers of aging, our results apply to non-molecular markers as well. Indeed, the most commonly used non-molecular markers of aging, such as grip strength or skin elasticity (50, 51), are non-causal.

Our results show that such physiological measurements, if sufficiently strongly correlated with mortality, can be used to identify causal loci of aging. However, a longitudinal study is required first to identify reliable physiological markers of mortality rates, whether molecular (52), physical (53, 54), or mental (55). What we have shown is that it is possible for a cross-sectional study to “piggy back” off markers identified from a longitudinal study to identify other, causative, loci of aging.

To better identify causal factors of aging we need a better understanding of what it means to be “aged”. Regression analyses using subjects’ chronological ages implicitly assume that “aging” is the process of having survived for a certain duration of time. Alternatively, “age” can be interpreted as morbidity, with associated mortality rate following a Gompertz curve. Ultimately, we are interested in the causal factors that prevent survival or decrease quality of life beyond a certain age. Our results confirm the critical importance of long-term longitudinal studies, whether directly conducted or indirectly incorporated, in finding both correlates and causes of aging.

## Author Contributions

Paul Nelson designed the methods, conducted the simulations, and wrote the manuscript. Daniel Promislow raised the initial questions regarding biomarkers, and contributed to the writing of the manuscript. Joanna Masel contributed to the methods and the writing of the manuscript.

This work was supported by the National Institute of Health (grant number K12GM000708); and the John Templeton Foundation (grant number 60814). Daniel Promislow was supported in part by grant DMS1561814 from the National Science Foundation, and grant AG049494 from the National Institute of Health.

